# A telomere-to-telomere genome of *Manilkara zapota* (Chiku) reveals the repeat-mediated genome evolution

**DOI:** 10.64898/2026.09.02.748860

**Authors:** Manohar S. Bisht, Mitali Singh, Martin Abraham Puthumana, Sonal Sharma, Vineet K. Sharma

## Abstract

*Manilkara zapota* (Chiku) is a tropical fruit tree in the Sapotaceae family. In this study, we assembled the first Telomere-to-telomere (T2T) genome assembly of *M. zapota*, spanning 1.6 Gb with high contiguity (N50 = 134 Mb). Comparative analysis revealed a very recent divergence and a shared WGD among Sapotaceae members. A newly constructed Ancestral Sapotaceae Karyotype (ASK) suggests a fusion-dominated chromosome structure in the family. The genome exhibits lineage-specific proliferation of transposable elements (TEs) accompanied by a recent burst of *LTR*-*Gypsy* elements (∼0.09 mya). Genome-wide methylation profiling revealed elevated CpG and CHG methylation levels and age-dependent reinforcement of TE silencing, suggesting strong epigenetic regulation that stabilises the repeat-rich genome while maintaining gene expression. Collectively, these findings advance our understanding of repeat-rich genome evolution in plants and provide a valuable framework for genomic-assisted breeding in *M. zapota*.

## Introduction

*Manilkara zapota* (2n=26), commonly known as Chiku, is a tropical evergreen tree from the Sapotaceae family of flowering plants, native to southern Mexico. It is widely distributed throughout America, the Caribbean, and Southeast Asia, and is heavily cultivated across India for its commercially important sweet fruit. The tree can grow up to about 20m in height, the leaves are elliptic to lanceolate, and the flowers are white, bisexual, and bell-shaped. The fruit is brown and oval, measuring about 5-10 cm (**Figure 1A**). Chiku fruit has a high sugar content (12-14% fructose and sucrose), substantial amounts of carbohydrates, protein, and minerals, and significant nutritional value (Rivas-Gastelum *et al*., 2023). In addition to its nutritional properties, *M. zapota* contains various bioactive compounds, including alkaloids, flavonoids, tannins, and saponins, which confer robust medicinal properties. Various parts of the plant, including leaves, fruit, bark and seeds, have been known to confer antimicrobial, antidiabetic, antioxidant, anti-inflammatory, and anticancer activities (Gam *et al*., 2024; Rivas-Gastelum *et al*., 2023).

**Figure 1:**
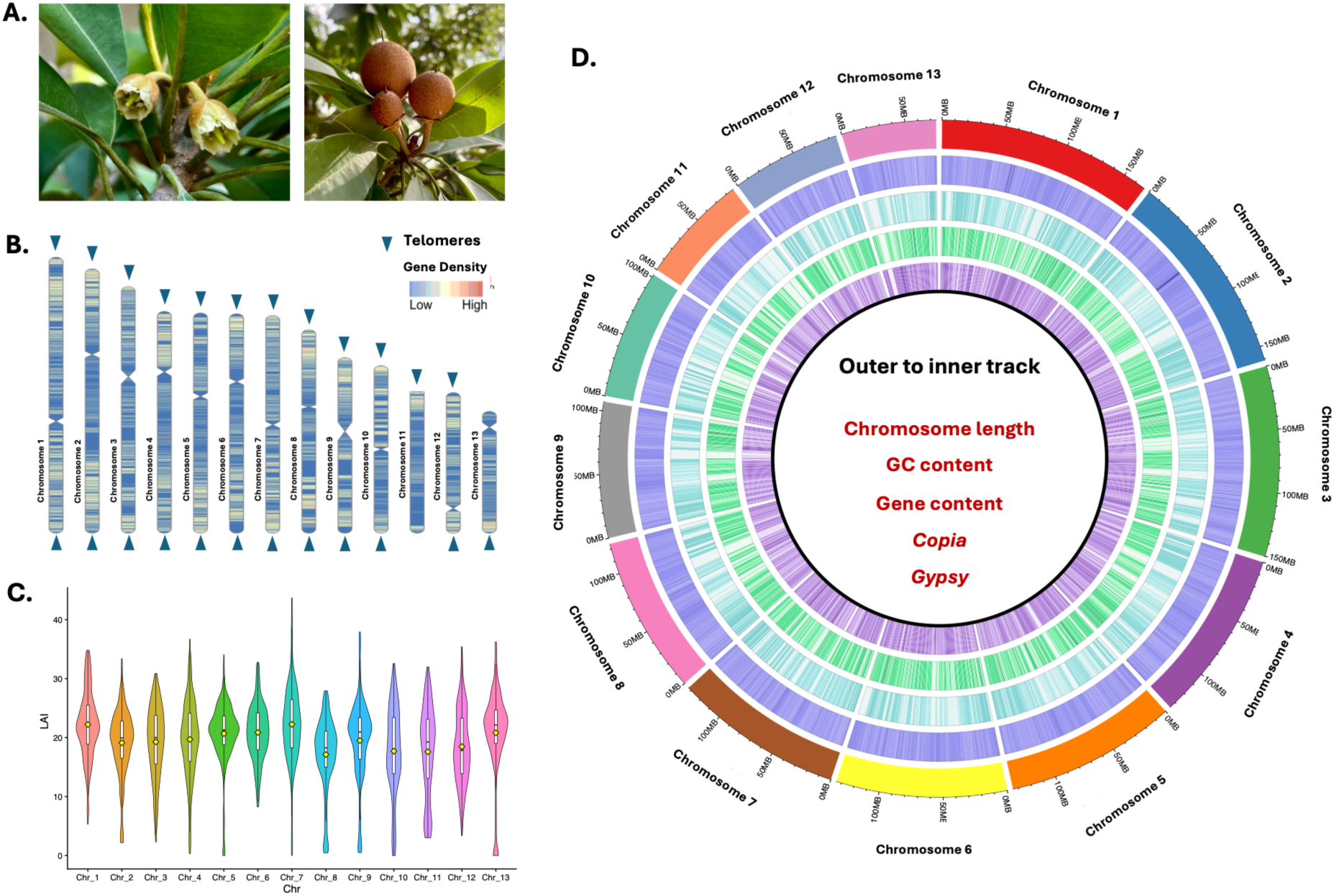
Morphology and genomic characteristics of *M. zapota*. **A.** Morphological characteristics of *M. zapota*, including flowers (left) and fruits (right). **B.** Gene density and the distribution of telomeres and centromeres. **C.** Distribution of LAI scores for chromosomes. **D.** Circos plot illustrating genome-wide features, including chromosome length (outermost track), GC content, gene density, and distribution of major transposable element classes (*Copia* and *Gypsy*) from outer to inner tracks. All distributions are drawn in a window size of 100 kb.

The emergence and diversification of specialised metabolic pathways in plants are frequently associated with structural variation in the genome, including gene duplication, transposition, and transposable element (TE)-mediated genome remodelling. Increasing evidence suggests that TEs not only contribute to genome expansion but can also facilitate metabolic innovation by generating novel regulatory variation, promoting gene family expansion, and reshaping the genomic architecture of biosynthetic pathways. However, the extent to which TE proliferation contributes to the evolution of specialised metabolism remains poorly understood, particularly in non-model plant species. Comparative analyses across closely related taxa provide a powerful framework to disentangle these processes. A robust understanding of TE-driven genome evolution requires the study of genomes in which phylogenetic distance is minimised and chromosomal organisation is well conserved, enabling direct comparisons of genomes that share regulatory backgrounds yet exhibit substantial differences in TE content. However, only a limited number of such studies are available (Buendía-Ávila *et al*., 2026; Igolkina *et al*., 2025; Quadrana *et al*., 2019; Wyler *et al*., 2020; Zemach *et al*., 2010). In this context, the Sapotaceae family represents such a framework, in which closely related species retain a conserved karyotypic structure yet exhibit pronounced differences in repeat content and genome size, thereby providing a unique opportunity to investigate how repeat expansion has influenced genome evolution.

This study reports the first high-quality T2T assembly of the highly repetitive genome of *M. zapota*. The comparative genomic analyses revealed the evolutionary history of Sapotaceae, including divergence patterns, a shared whole-genome duplication event, and reconstruction of the ancestral karyotype. The study also provides the first comparative insights into the landscape and role of repetitive elements in genome expansion and regulation. Together, by linking repeat expansion, epigenetic regulation, and metabolic innovation, our study advances understanding of repeat-rich genome evolution in plants.

## Results

### Assembly, assessment and annotation of the *M. zapota* genome

The genome assembly was constructed using the Oxford Nanopore long reads (115 Gb) and Illumina short reads (220 Gb) data. Genome survey analysis using GenomeScope2 (K=31) estimated the haploid genome size to be approximately 1.41 Gb with heterozygosity of ∼1.7%. The primary contig assembly from HiFiasm was 1.63Gb in length with a N50 of 77.3 Mb. With the integration of HiC reads, a high N50 value of 134 Mb could be achieved. Further, it helped in anchoring the contig-level genome to 13 superscaffolds (chromosomes) at a ∼97% anchor rate with chromosomes ranging from 168.4 Mb to 74.3 Mb. Furthermore, a total of 24 telomeres and 12 centromeric sequences were identified from the genome assembly (**Figure 1B**).

Multiple parameters were used to assess assembly quality, including BUSCO analysis, which reported a completeness score of 99.57%. The average LAI of the constructed chromosomes was 19.33, close to the “gold standard” (**Figure 1C**). The assessment using Merqury reported a quality value (QV) score of 55.02 across the 13 chromosomes, corresponding to an estimated single-base accuracy of 99.999%. In addition, 99.94% of Illumina short reads and 97.97% of Nanopore long reads could be mapped to the final genome assembly. These quality metrics collectively suggest an excellent-quality genome assembly of *M. zapota* (**Figure 1D**).

A total of 1.28 Gb (78.61% of the genome) bases in the *M. zapota* genome were constituted of repeats. Of these, 53.09% was contributed by Retroelements and 14.60% by DNA transposons. A total of 35,612 coding genes could be identified (longest isoforms only) using the *ab-initio* and evidence-based modes on the repeat-masked genome. These high-confidence coding genes showed a BUSCO completeness of 98.9%, and 95% of these genes could be functionally annotated.

### Phylogenetic relationship, evolutionary history, and gene family evolution

To understand the phylogenetic relationship among the Sapotaceae species, a time-calibrated maximum likelihood (ML) tree based on 1,084 one-to-one fuzzy orthogroups was constructed. The analysis revealed that the Sapotaceae family diverged from other Ericales order species at around 52 million years ago (mya). Within Sapotaceae, a recent divergence among the selected species was observed, where *Sideroxylon spinosum* was the first to diverge (23 mya), followed by *Synsepalum dulcificum* (21 mya). *M. zapota* was found to be recently diverged from its sister clade, *V.* pardoxa and *M. longifolia* (**Figure 2A**). To further examine the recent divergence among Sapotaceae members, the Ks analysis of orthologous genes was performed, which revealed a corroborated divergence pattern consistent with that observed in the time-calibrated phylogeny (**Figure 2B(i)**). The WGD analysis using paralogous genes revealed that all selected Sapotaceae members shared a duplication event around Ks ∼ 0.55 before diverging, and no additional independent WGD events occurred in the selected Sapotaceae species (**Figure 2B(ii)**). In addition, strong syntenic conservation and 1:1 syntenic depth were observed among Sapotaceae species, providing further evidence of recent divergence and shared WGD (**Figure 2C** and **S2**).

**Figure 2:**
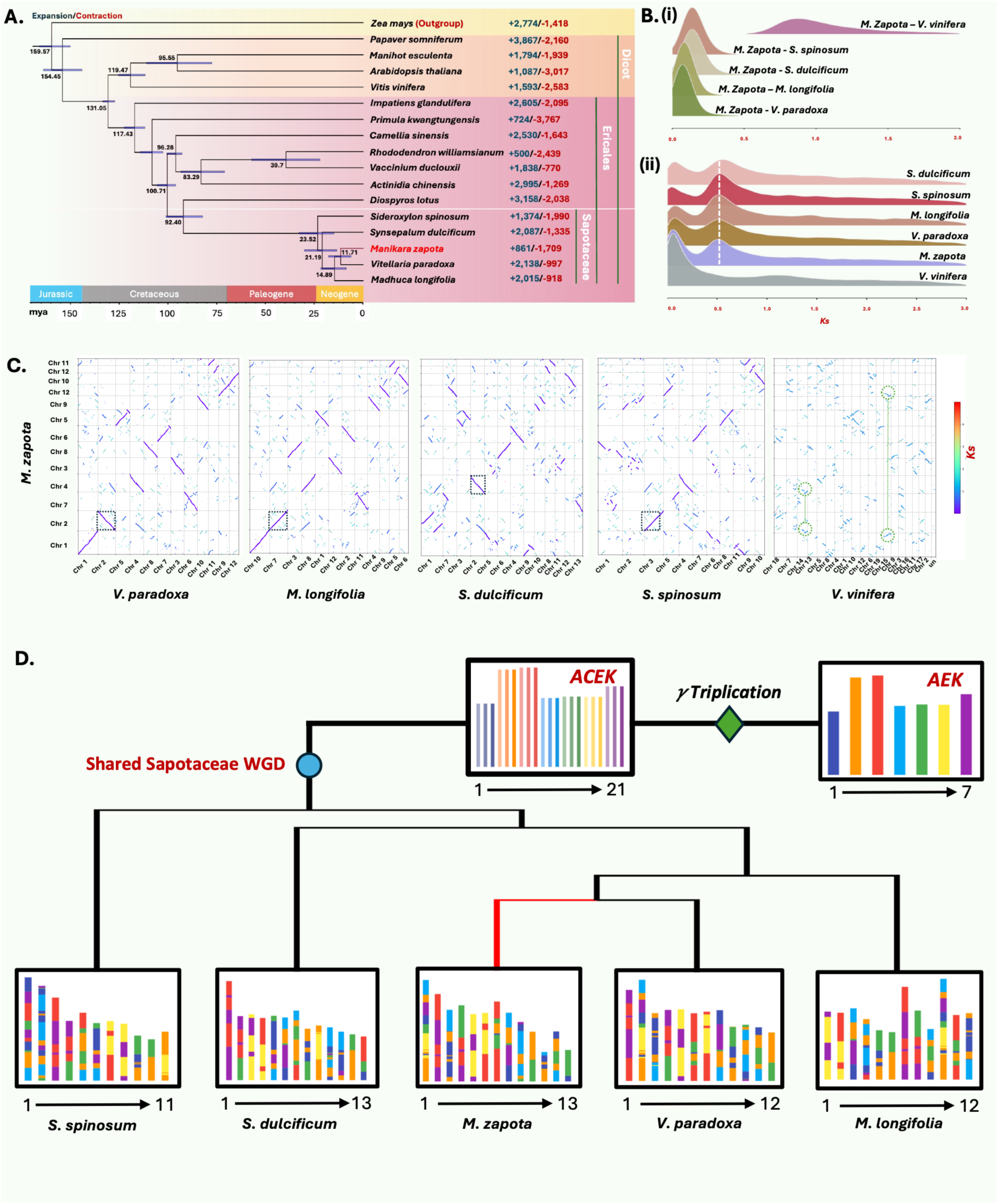
Comparative genome evolution in Sapotaceae. **A.** Phylogenetic tree showing divergence times and gene family expansion/contraction across representative species, with *Zea mays* as the outgroup. Numbers on branches indicate expanded and contracted gene families. **B.** Ks (synonymous substitution rate) distributions illustrating duplication events: **(i)** pairwise ortholog comparisons between *M. zapota* and other species, and **(ii)** species-specific Ks distributions. Dotted lines highlight lineage-specific duplication patterns. **C.** Synteny dot plots comparing *M. zapota* with other Sapotaceae species and *V. vinifera*, revealing conserved collinearity and duplication signals across chromosomes. Dotted boxes represent 1:1 synteny. **D.** Schematic representation of reconstructed karyotypes of Sapotaceae species highlighting the shared Sapotaceae whole-genome duplication event and the ancestral γ (gamma) triplication, along with inferred chromosomal rearrangements and karyotype evolution across species.

To understand the karyotype evolution of Sapotaceae species, the Ancestral Sapotaceae Karyotype (ASK) with 13 protochromosomes was constructed using the shared intact chromosome-level collinear blocks among the early-diverging members of the Sapotaceae. The mapping of ASK against the five Sapotaceae species revealed lineage-specific chromosomal rearrangements that led to their modern karyotype structures. In *M. zapota*, Chr1 and Chr3 were formed from ASK5, ASK9 and ASK11 through EEJ (end-to-end joining), NCF (nested chromosome fusion), fission events, and chromosomal inversions. Whereas Chr7 and Chr12 are formed by the RTA (reciprocal translocated chromosome arms) event between ASK7 and ASK13. Further, *M. longifolia* and *V. paradoxa* also shared a similar chromosomal evolutionary trajectory to *M. zapota*. The shared chromosomal fusion events in these lineages have resulted in 12 chromosomes. However, in *M. zapota*, ASK8 underwent a separate fission event to produce Chr11 and Chr13. Despite undergoing WGD after WGT (triplication), the ASK has a chromosome number of 13, suggesting that Sapotaceae members underwent additional fusions, as depicted by the karyotype projection of five Sapotaceae species from the AEK (**Figure 3D**). However, the additional evidence from the genome assemblies of this family will further refine the inferred ancestral karyotype of Sapotaceae.

**Figure 3:**
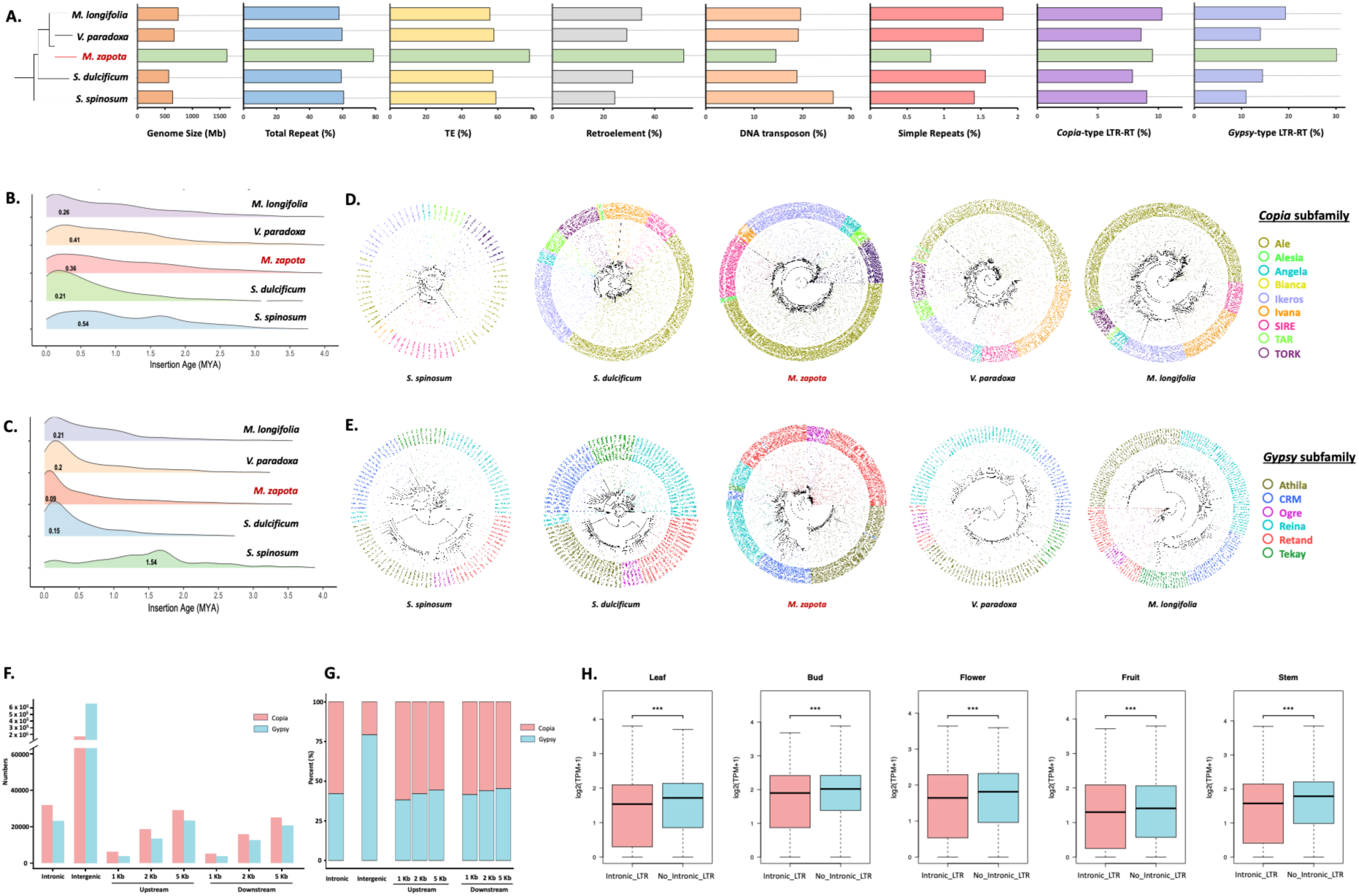
Comparative TE landscape. **A.** Comparative analysis of genome size and repeat composition across Sapotaceae species, including total repeat content, TEs, retroelements, DNA transposons, simple repeats, and proportions of *Copia* and *Gypsy* LTR retrotransposons. **B-C.** Insertion age distributions of *Copia* and *Gypsy* respectively. **D-E.** phylogenetic relationships between *Copia* and *Gypsy* subfamily, respectively. **F–G.** Distribution and proportion of *Copia* and *Gypsy* elements across genomic regions, including intronic, intergenic, and flanking regions (upstream and downstream), indicating preferential accumulation in specific genomic compartments. **H.** Comparison of gene expression levels (log₂(TPM+1)) between genes with and without intronic LTR insertions. Significant differences between groups are indicated by *** (Wilcoxon rank-sum test; p < 0.001). *n=3* for bud and flower, *n=2* leaf and stem, and *n=1* for fruit.

The analysis of gene family evolution in *M. zapota* revealed expansion in 861 gene families and contraction in 1,709 families (**Figure 2A**). Functional enrichment analysis revealed that the expanded gene families were predominantly associated with secondary metabolism and core cellular processes. Notably, pathways related to terpenoid biosynthesis, including monoterpenoid and sesquiterpenoid/triterpenoid biosynthesis, were significantly enriched, along with phenylpropanoid and flavonoid biosynthesis pathways.

### Burst of repeat sequences in the *M. zapota* genome

The *M. zapota* genome represents an extreme case of recent, repeat-driven expansion within Sapotaceae. The genome of *M. zapota* was found to be twice the size compared with other genomes of the Sapotaceae family (available at the time of study). A detailed comparative gene-architecture analysis across Sapotaceae genomes revealed that coding sequence space remains relatively conserved across the family, with total exon lengths ranging from 32 to 42 Mb. In contrast, substantial variation was observed in intronic and intergenic regions. The *M. zapota* genome exhibited the largest gene span (212 Mb) and extensive intergenic expansion (∼1.39 Gb), accompanied by the longest average intron length (1,357 bp) compared to other members of the Sapotaceae family. Conversely, the *S. dulcificum* genome exhibited the most compact gene architecture, with shorter introns and intergenic regions (∼446 Mb). These findings indicated that genome size differences among Sapotaceae species were primarily driven by intron and intergenic expansion, likely resulting from lineage-specific transposable element (TE) proliferation rather than gene gain.

The repeat analysis also supported the above observations. An exceptional accumulation of transposable elements (TEs) accounting for ∼78% of the *M. zapota* genome was substantially higher than the other family members (57–61%). Notably, the TE class in *M. zapota* was dominated by retrotransposons, whereas DNA transposons were comparatively less abundant than in other Sapotaceae genomes. Among retrotransposons, long terminal repeat (LTR) elements were particularly enriched, with *LTR-Gypsy* showing a pronounced predominance compared to *LTR-Copia*, suggesting a lineage-specific expansion of *Gypsy*-type retrotransposons (**Figure 3A**). Insertion time analysis further revealed a recent and asymmetric burst of LTR activity, with Copia elements expanding earlier (∼0.36 mya), and a more recent Gypsy burst (∼0.09 mya) (**Figure 3B** and **C**). These findings identify a recent, Gypsy-driven expansion as the primary force underlying genome enlargement in *M. zapota*.

Phylogenetic classification of intact LTR elements uncovered distinct subfamily-level expansions, with *Copia* elements dominated by Ale, and *Gypsy* elements by Athila lineages. Notably, several subfamilies, including SIRE and Ikeros (*Copia*) and Retand and CRM (*Gypsy*), exhibited higher amplification specifically in *M. zapota*, whereas other elements, such as Tekay, remained comparatively constrained. This pattern highlights highly selective, lineage-specific TE expansions rather than uniform proliferation across families (**Figure 3D** and **E**).

At the genomic scale, LTR elements display strong spatial organisation, with intergenic regions heavily enriched in *Gypsy* elements (>80%), while intronic regions show relatively higher Copia representation. TE density increases with distance from genes, reflecting progressive accumulation in gene-distal regions (**Figure 3F** and **G**). Further, genes with intronic LTR insertions (8,897) showed modest but reduced expression compared to length-matched control genes lacking LTR insertions (26,716) across all five tissues (p <0.001) (**Figure 3H**). This near-neutral transcriptional impact likely contributes to the evolutionary persistence of these insertions, providing a substrate for potential regulatory co-option over longer evolutionary timescales.

### Genome-wide methylation profile and expression regulation in *M. zapota*

The repeat-rich genome of *M. zapota* revealed a higher genome-wide DNA methylation. Comparative analysis with the moderately repetitive genome of *M. longifolia* revealed that *M. zapota* exhibited substantially higher methylation across all sequence contexts, with CpG (71.46%), CHG (55.06%), and CHH (10.08%) levels consistently exceeding those observed in *M. longifolia*. This global increase indicates enhanced epigenetic suppression associated with large-scale TE accumulation.

At the gene level, methylation profiles revealed a conserved regulatory architecture, with reduced methylation at transcription start and end sites (TSS/TES) and elevated methylation across gene bodies and flanking regions in both species. However, *M. zapota* showed a markedly stronger separation between CpG and CHG methylation than *M. longifolia*, reflecting greater maintenance of symmetric methylation and more robust transposon silencing (**Figure 4A(i)**). Further, exonic regions exhibited relatively lower methylation across all three sequence contexts in both genomes, whereas intronic regions showed higher methylation levels, which were significantly enriched in *M. zapota* compared with *M. longifolia*, indicating increased targeting of intragenic repetitive elements and TE-derived insertions in the expanded genome and also suggesting that the high methylation in gene-body regions was caused by repeat-rich introns (**Figure 4A (ii-iii)**). Consistent with this, repeat regions including *Copia* and *Gypsy* LTR retrotransposons were strongly hypermethylated in *M. zapota*, particularly in symmetric contexts (CpG and CHG), whereas CHH differences were comparatively less. This pattern highlights a dominant role for maintenance methylation in stabilising the expanded TE landscape (**Figure 4A (iv-vi)**).

**Figure 4:**
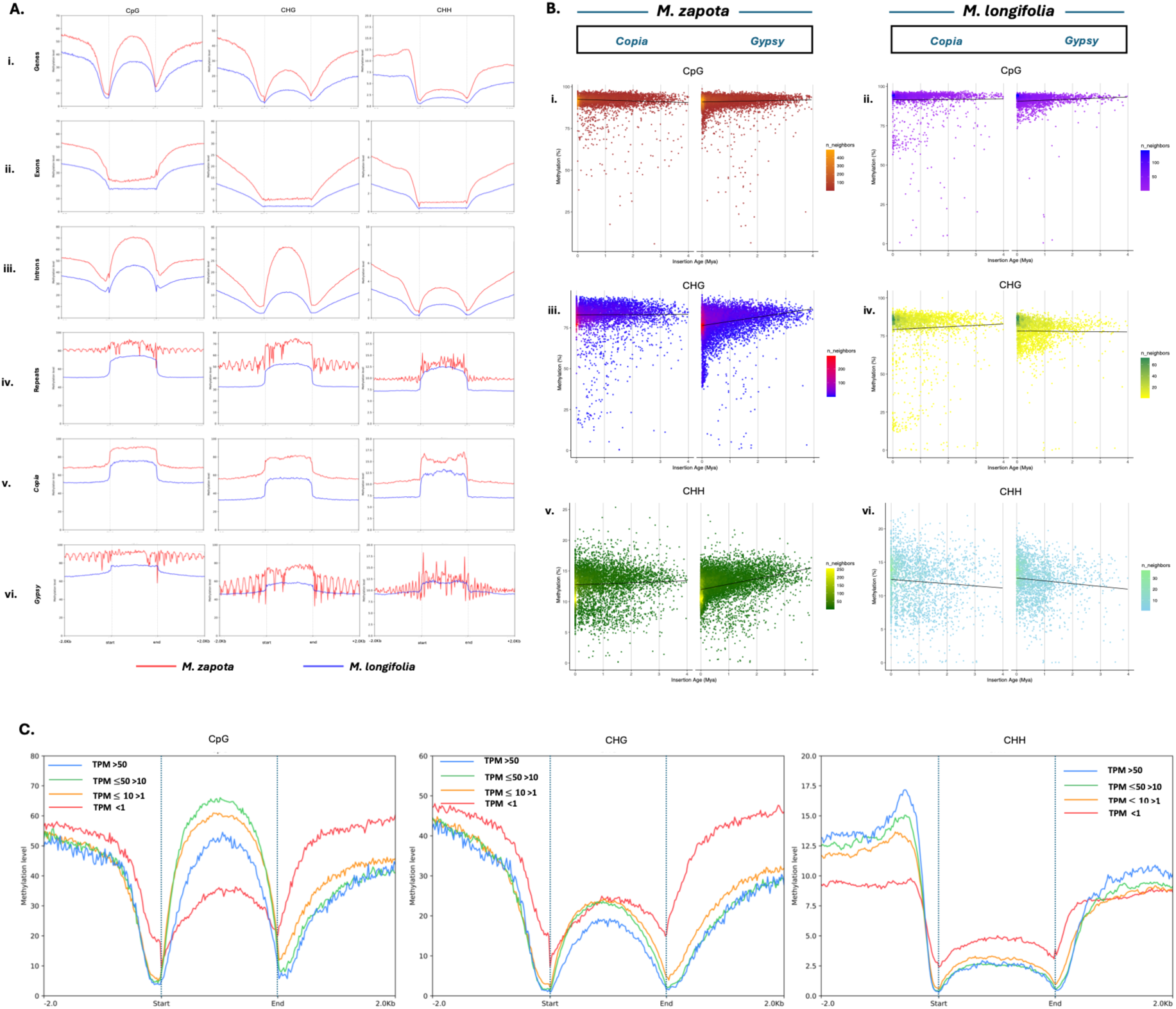
Genome-wide DNA methylation analysis in *M. zapota*. **A.** Average DNA methylation levels (%) in CG, CHG and CHH sequence contexts across genomic features (genes, exons, introns, repeats, *Copia*, and *Gypsy* elements) in *M. zapota* and *M. longifolia*. **B.** Relationship between transposable element insertion age and DNA methylation levels for *Copia* and *Gypsy* elements, illustrating age-dependent methylation dynamics across CpG, CHG, and CHH contexts in *M. zapota* and *M. longifolia* species. **C.** Comparison of DNA methylation profiles and expression levels between four expression categories based on TPM.

To examine the relationship between LTR insertion age and epigenetic regulation, methylation levels were analysed in correlation with insertion age between *M. zapota* and *M. longifolia*. In both species, CG methylation remained consistently high across all insertion age classes, indicating stable maintenance methylation and long-term silencing of transposable elements. However, *M. zapota* has a higher density and spreads at a young age (< 0.2 mya), indicating the recent LTR burst (**Figure 4B(i)**) compared to *M. longifolia*. CHG methylation of *Copia* elements increased progressively with insertion age in both species. Further, a markedly stronger increased trend was observed among Gypsy elements with insertion age in *M. zapota* compared to *M. longifolia*. This pattern indicates progressive reinforcement of heterochromatic silencing following retrotransposon insertion and suggests a reduced turnover of older elements in the repeat-rich genome of M. *zapota*. In contrast, the weaker age-dependent CHG increase observed in *M. longifolia* suggests a more dynamic transposon removal and reduced long-term retention of LTR retrotransposons, contributing to its comparatively smaller genome size (**Figure 4B (ii)**). In contrast to the canonical decline of CHH methylation with transposable element age observed in *M. longifolia*, Gypsy elements in *M. zapota* exhibited an increase in CHH methylation with insertion age LTR retrotransposons (**Figure 4B (iii)**). This unexpected pattern suggests persistent RNA-directed DNA methylation targeting of older retrotransposons in the repeat-rich genome of *M. zapota*. Such long-term reinforcement of CHH methylation is consistent with the accumulation of *Gypsy* elements within heterochromatic regions and likely contributes to the stabilisation and retention of transposable elements in the expanded genome of *M. zapota* species.

Given the globally elevated cytosine methylation observed in *M. zapota* across CG, CHG, and CHH contexts, the relationship between methylation patterns and gene expression levels was investigated. Genes were classified into four expression categories based on TPM values (see Methods). A clear opposite relationship was observed between methylation in gene-flanking regions and transcriptional activity, particularly around TSS and TES. The highly expressed genes (TPM > 50) exhibited substantially reduced methylation compared with moderately and weakly expressed genes. In contrast, CpG methylation within gene bodies showed a positive association with transcriptional activity, consistent with the conserved pattern of gene-body methylation observed in many plant species (**Figure 4C**). Together, these findings indicate that role of DNA methylation in shaping transcriptional regulation in *M. zapota,* where methylation in promoter-proximal and downstream regulatory regions results in transcriptional repression, whereas gene-body CG methylation contributes to transcriptional stability.

### Genome-wide characterisation of transposed duplicated genes in *M. zapota*

A total of 31,044 duplicated genes were identified in the M. zapota genome. These genes were further divided into five categories: proximal duplication (PD) genes (4,182), tandem duplication (TD) genes (3,919), dispersed duplication (DSD) genes (3,901), transposed duplication (TRD) genes (5,736), and WGD genes (13,306) (**Figure 5A**). Further analysis of duplicate gene types revealed that PD and TD exhibited higher Ka/Ks ratios than DSD, TRD and WGD, suggesting relatively relaxed purifying selection acting on PD and TD duplicates, compared to stronger purifying selection observed in DSD, TRD and WGD duplicates (**Figure 5B**). This pattern may be explained by the fact that WGD, DSD, and TRD duplicates tend to undergo exponential decay over evolutionary time, whereas PD and TD continuously contribute new gene copies to the genome (Qiao *et al*., 2019). Among the duplicated gene categories identified in Sapotaceae genomes, TRD genes represented the second most abundant duplication type, after WGD genes. However, because of the shared WGD event with selected Sapotaceae species, the functional contribution of WGD-derived genes was not further investigated. Instead, a detailed analysis of TRD genes was performed. KEGG pathway enrichment analysis of TRD gene pairs revealed significant enrichment of transposed genes in key primary metabolic pathways, including carbon metabolism, glycolysis, and starch and sucrose metabolism, as well as secondary metabolic pathways such as flavonoid and phenylpropanoid biosynthesis, compared to the parental copies (**Figure 5C**).

**Figure 5:**
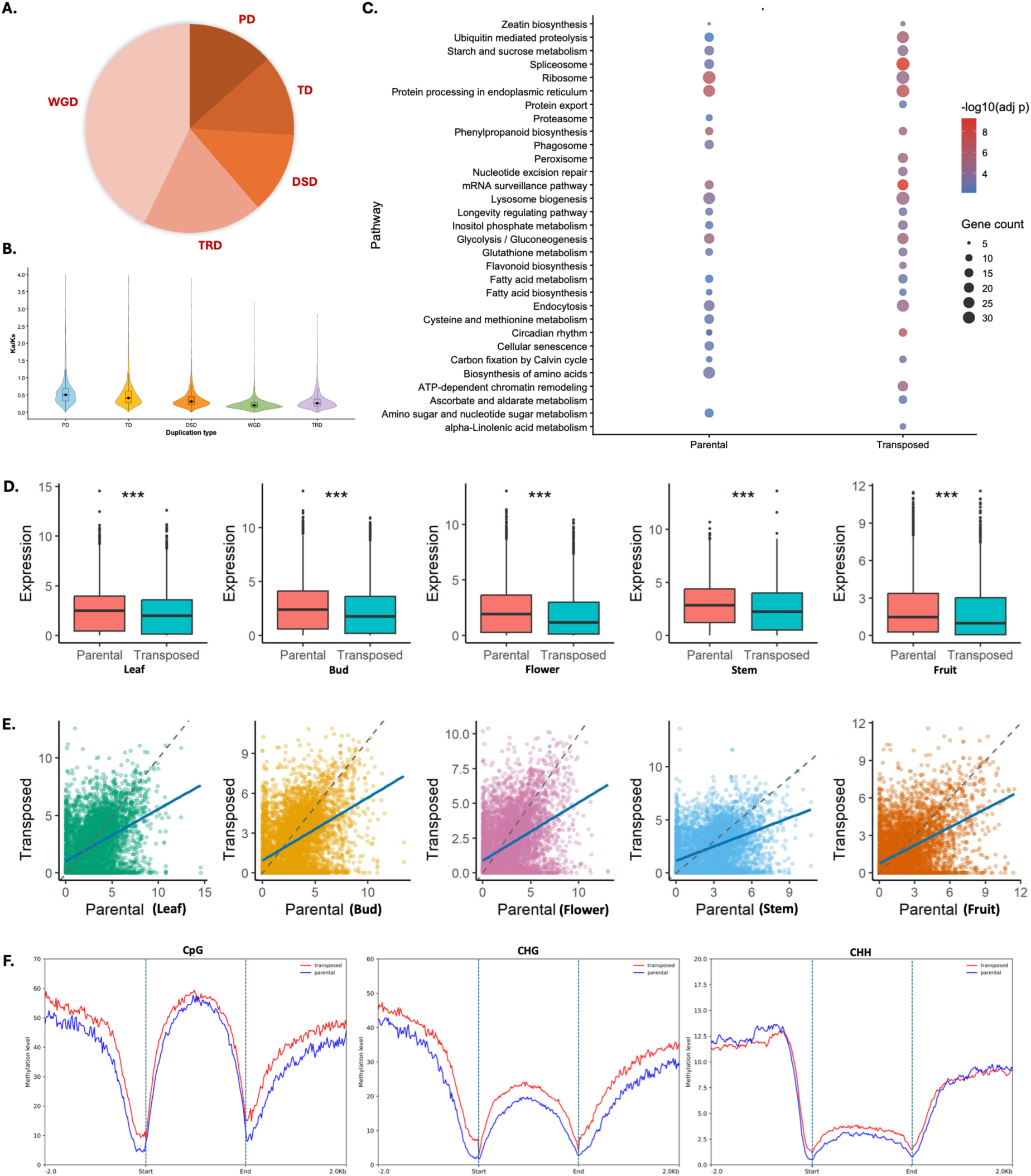
Functional enrichment, expression divergence, and methylation landscape of transposed duplicate genes in *M. zapota*. **A.** Distribution of duplicate genes across different duplication classes, including proximal duplicated (PD), tandem duplicated (TD), dispersed duplicated (DSD), transposed duplicated (TRD), and whole-genome duplicated (WGD) genes. **B.** Distribution of Ka/Ks ratios among different duplication classes. **C.** KEGG pathway enrichment analysis of parental and transposed duplicate genes, highlighting the functional categories significantly enriched in each gene set. **D.** Comparison of expression levels between parental and transposed duplicate genes across five tissues (leaf, bud, flower, stem, and fruit). Box plots represent the distribution of normalised expression values, and significant differences between parental and transposed copies are indicated by asterisks (Wilcoxon rank-sum test; ***, P < 0.001). *n=3* for bud and flower, *n=2* leaf and stem, and *n=1* for fruit. **E.** Correlation of expression levels between parental and transposed duplicate genes across different tissues. Each point represents a duplicate gene pair, and the solid blue line denotes the linear regression fit. **F.** Average DNA methylation profiles of parental and transposed duplicate genes across gene bodies and 2-kb upstream and downstream flanking regions in the CpG, CHG, and CHH sequence contexts. Methylation levels are shown relative to the transcription start site (TSS) and transcription end site (TES), with dashed vertical lines indicating gene boundaries.

To further assess the evolution of transposed genes relative to their parental genes, expression levels of transposed and parental genes were compared. Notably, the parental gene was expressed at significantly higher levels than the transposed gene across all examined tissues (p < 0.001). Furthermore, a weak correlation in expression patterns was observed between the two gene types (**Figure 5D** and **E**). In addition, DNA methylation profiling revealed consistently higher methylation levels in transposed gene copies compared to their parental counterparts across all three contexts (CpG, CHG, and CHH). This difference was particularly pronounced in the gene body and flanking regions. Elevated CHG and CHH methylation in transposed genes suggests their association with transposable element–rich genomic environments and potential targeting by silencing pathways (**Figure 5F**). Together, these results suggest that transposed duplicates were not functionally lost but underwent regulatory modulation and partial divergence, contributing to genome evolution through a balance of functional retention and diversification.

## Discussion

In this study, we have constructed a high-quality T2T genome of *M. zapota* (Sapotaceae), a commercially important fruit crop with industrial applications. The high repetitive content and heterozygosity of the genome posed challenges for the assembly, however, the high N50, BUSCO, LAI and the identification of telomeric and centromeric sequences achieved for this assembly attest to the genome quality.

The evolutionary and phylogenomic analyses carried out in this study revealed that Sapotaceae members diverged from other Ericales around 52 mya, with a very recent divergence and a shared WGD event between the Sapotaceae species. In addition, the reconstruction of the ancestral Sapotaceae karyotype suggested that chromosome evolution within the family has been largely driven by fusion-dominated structural rearrangements following ancient polyploidization events. The conservation of the ancestral chromosome number in *M. zapota*, despite evidence of lineage-specific rearrangements, indicates a relatively stable karyotype trajectory. These findings provide new insights into the structural evolution of Sapotaceae genomes and establish a framework for future comparative genomic analyses as additional chromosome-level assemblies become available.

In the Sapotaceae family, *M. zapota* genome showed a lineage-specific genome expansion. The recent burst of transposons has led to the expansion of intronic and intergenic regions of the genome. Further, the disproportionate accumulation and burst of *LTR-Gypsy* (0.09 mya) elements relative to *LTR-Copia* (0.36 mya) in *M. zapota* indicates that genome expansion in this lineage is not only recent but ongoing, showing a punctuated retrotransposon activity similar to other studies where rapid post-speciation TE amplification accounts for the bulk of interspecific genome size differences (Palmer *et al*., 2012; Pulido and Casacuberta, 2023). Notably, a preferential enrichment of *Gypsy* elements in intergenic and *Copia* elements in intronic regions further suggests that the two superfamilies occupy distinct genomic niches within *M. zapota*, potentially reflecting differences in their integration site preferences mediated by their respective integrase chromodomains (Pereira, 2004; Zhang *et al*., 2020).

Our comparative whole-genome methylome analysis supports the view that TE content and distribution are major parameters shaping the divergent epigenomes between *M. zapota* and *M. longifolia*. The abundance of symmetric methylation (CpG and CHG) and increased intragenic methylation in *M. zapota* suggests a strong maintenance of heterochromatin and stable repression of TE-derived insertions, contributing to genome expansion and structural stability. Further, the age-dependent methylation patterns of retrotransposons indicate a prolonged retention and reinforcement of silencing in older, particularly *Gypsy* elements, in contrast with the more dynamic turnover observed in *M. longifolia*. In addition, at the gene level, the observed contrasting relationship between methylation and expression highlights the dual and context-dependent role of DNA methylation in transcriptional regulation.

In conclusion, this study presents the first high-quality T2T genome assembly of the commercially important fruit crop *M. zapota*. Together, by linking repeat expansion, epigenetic regulation, and metabolic innovation, our study advances understanding of how plant genomes evolve and generate phenotypic novelty. The comprehensive genomic and evolutionary insights into its repeat-rich genome dynamics and epigenetic regulation provide a valuable foundation for future investigations into repeat-driven genome evolution and metabolic innovation. These findings will further facilitate trait improvement and biotechnological applications in *M. zapota* and other tropical fruit crops.

## Methods

### DNA and RNA extraction and sequencing

DNA was extracted from the leaves of the plant growing in the IISER Bhopal campus (23.29 °N 77.27 ° E). CTAB lysis buffer (100mM Tris-Cl, 2% CTAB, 2M NaCl, 1% PEG8000, and 20mM EDTA, pH=9) was used for lysis, followed by chloroform: isoamyl extraction and precipitation of DNA in aqueous extract with isopropanol. The DNA pellet was eluted in Tris-Cl and stored. The quality of the extracted DNA was checked on NanoDrop™ 8000 Spectrophotometer (ThermoFisher Scientific, USA) and quantification was done on Qubit 2.0 fluorometer. For short read sequencing the library preparation was carried out using KAPA HyperPlus Kit (Roche) and sequenced on the Illumina NovaSeq 6000 platform. For the long read sequencing the extracted DNA was purified with the Genomic-tip 20/G (Qiagen). 10 micrograms of the DNA was added to the G2 buffer with 4 ul RNaseA (PureLinkTM, Invitrogen) and heated at 56 °C for 15 min. The solution was then loaded onto the equilibrated genomic tip and was allowed to pass through, followed by 3X washing with Buffer QC, and elution in Buffer QF. The DNA was precipitated by adding 0.7x ice-cold isopropanol to the eluate, followed by pelleting of the DNA, ethanol washing of the pellet, and elution in Tris-Cl (pH 8.5). The library was prepared using the ligation kit SQK-LSK114 and loaded onto. A total of two runs on the P2 Solo generated more than 100 GB of data. The first run was with ∼60 GB, N50 13.63kb, and the second run was with ∼40 GB of data, with read N50 of 26.82kb. For the transcriptome analysis, multiple tissues (leaf, stem, flower buds, open flower and fruit) were collected. RNA extraction was performed following the method described in Singh et al., 2026 (Singh *et al*., 2026). Extracted RNA was purified using the Qiagen RNeasy mini kit spin column following the manufacturer’s protocol, with on-column DNA digestion using Ambion TM DNase I (Invitrogen) prior to library preparation and sequencing.

### Hi-C library preparation and sequencing

The Hi-C library was prepared with young and freshly collected leaves of the plant. The tissue was first fixed in 1% formaldehyde, followed by nuclei isolation and further processing using the EpiTect Hi-C kit (Qiagen), following the manufacturer’s protocol. The sequencing was done on the Illumina NovaSeq 6000 platform.

### Genome assembly and quality assessment

The genome size was predicted from filtered Illumina short reads using k-mer analysis with Jellyfish v2.2.8 (Marçais and Kingsford, 2011) and visualised using GenomScope2 (Ranallo-Benavidez *et al*., 2020). HiFiasm v0.14-r312 (Cheng *et al*., 2021) was used to construct a contig-level assembly by combining ONT long reads and HiC data (-HiC mode) with “–l 3” and “-s 0.5” parameters. The primary assembly was further scaffolded to construct a pseudo-chromosome assembly using YAHS v 1.2.2 (Zhou *et al*., 2023) with the “--no-contig-ec” option. The HiC matrix was then created using Juicer v1.2.2 (Durand, Shamim, *et al*., 2016), and manually curated using Juicebox (Durand, Robinson, *et al*., 2016). LR_Gapcloser was used in 5 iterations to fill gaps in the final assembly (Bisht *et al*., 2024; Xu *et al*., 2018). The quality and completeness of the final genome assembly were evaluated using multiple benchmarking approaches, including QUAST v5.2.2 (Gurevich *et al*., 2013), BUSCO (embryophyta_odb10) (Huang and Li, 2023; Manni *et al*., 2021), and Merqury v1.3 (Rhie *et al*., 2020). Assembly accuracy and coverage consistency were further assessed by mapping genomic long and short reads back to the assembly using minimap2 v2.17-r941 (Li, 2018) and bwa-mem 0.7.17-r1188 (Li and Durbin, 2009), respectively. Additionally, telomeric and centromeric sequences were predicted using quarTeT v1.2.5 (Lin *et al*., 2023) and CentIER v1.1 (Xu *et al*., 2024), respectively.

### Gene set construction and functional annotation

The genome assembly from the previous step was used to construct a repeat library with EDTA v2.1.3 (Ou *et al*., 2019), and the repeats were soft-masked with RepeatMasker v4.1.2 (http://www.repeatmasker.org). The protein-coding genes were then annotated using the BRAKER3 pipeline (Gabriel *et al*., 2024), integrating RNA-seq data from different tissues and protein sequences from other Sapotaceae species as evidence. The obtained gene set was then filtered to retain only the longest isoforms per gene using GFFread v0.12.1 (Pertea and Pertea, 2020). Further, BUSCO was used to assess the completeness of the constructed gene set, along with various publicly available databases for functional annotation.

### Phylogenetic and gene family evolution analysis

Protein sequences from 15 species (including five Sapotaceae species) were used to construct the phylogeny. Orthogroups were created using Orthofinder v2.5.4 (Laetsch and Blaxter, 2017), and fuzzy one-to-one orthogroups containing the protein sequences were extracted using KinFin v1.2 (Emms and Kelly, 2019), which were further aligned by using MAFFT v7.467 (Katoh *et al*., 2019). The obtained multiple sequence alignments were concatenated and processed to remove empty sites using BeforPhylo v0.9.0 (https://github.com/qiyunzhu/BeforePhylo). RAxML v8.2.12 (Stamatakis, 2014) was used to construct a maximum-likelihood species phylogenetic tree using the ‘PROTGAMMAAUTO’ amino acid substitution model, with 100 bootstrap replicates (Chakraborty *et al*., 2021). Further, the divergence time was estimated using the MCMCtree implemented in PAML v4.10.6 (Yang, 2007) using six calibration points.

CAFE v5 (Mendes *et al*., 2021) was used to analyse expanded and contracted gene families. At first, all vs all BLASTP was performed to cluster and filter gene families. Then a two-lambda model was used to filter gene families output, and the ultrametric species tree was used to analyse the expansion and contraction of gene families (Bisht *et al*., 2025; Puthumana *et al*., 2025). The KEGG enrichment of expanded gene families was performed using clusterProfiler v4.18.0 (Yu *et al*., 2012).

### WGD analysis and the evolutionary history of Sapotaceae

To examine the WGD history of Sapotaceae species, Ks distributions of orthologous and paralogous genes between and within the species and *V. vinifera* were plotted using wgd v2.23 (Chen *et al*., 2024). Further, using the wgd viz function with mixture modelling parameters, the distribution of the substitution rate-aware corrected ortholog Ks was plotted.

To construct the Ancestral Sapotaceae Karyotype (ASK) and infer the karyotype evolutionary history of Sapotaceae, five available high-quality genomes from the Sapotaceae family were selected. The WGDI pipeline was implemented to identify ancestral chromosomes and reconstruct the ancestral karyotype of Sapotaceae (Sun *et al*., 2022). At first, pairwise comparisons of protein sequences from the genomes of all five species were performed using BLASTP (e-value 1e-3), and collinearity dot plots were generated with the “-d” parameter. Further, from these collinearity dot plots, the chromosome-level collinear blocks identified from the early-diverging members of the Sapotaceae, i.e., Argan vs Miracle, were considered as protochromosomes, and ASK was constructed using the “-ak’ parameter. The resulting ASK was then mapped to the five Sapotaceae species using the “-km” parameter to examine and visualise the chromosomal rearrangements. Finally, the published Ancestral Eudicot Karyotype (AEK) (Wang *et al*., 2022) was utilised to understand the karyotype projections of the reconstructed ASK and each of the five extant species using the “-k” parameter of the WGDI toolkit.

### Repeat and whole genome-wide methylation analysis

To perform a comparative repeat analysis among the five Sapotaceae members, the EDTA pipeline was used to identify all TEs. The insertion time for Copia and *Gypsy* elements was calculated using the intermediate files of the EDTA pipeline. Furthermore, the undamaged LTRs of Copia-type and *Gypsy*-type were further classified into more specific clades using TEsorter v1.3 (Zhang *et al*., 2022) software (https://github.com/zhangrengang/TEsorter/). The Phylogenetic trees for *Copia*-like and *Gypsy*-like LTR-RTs were constructed using IQtree v2.0.3 (Minh *et al*., 2020) software (http://www.iqtree.org/) using the protein sequences of the domain.

For identifying modified bases (5-Methylcytosine) in all contexts (CpG, CHG, and CHH) from ONT data, Dorado (v1.1.1) (https://github.com/nanoporetech/dorado) with the modification model “5mC_5hmC” was used against the reference genome assembly of *M. zapota* and *M. longifolia* to construct their corresponding modBAM files, respectively. The Modkit tool v0.6.1 (https://github.com/nanoporetech/modkit) was used to filter out base modifications of cytosine in each context, generate corresponding bedMethyl files, and convert them to bigWig files for visualisation in a genome browser. The deepTools suite v3.5.6 (Ramírez *et al*., 2014) was used to plot methylation profiles and generate heatmaps to understand genome-wide methylation patterns across multiple contexts, including repeat elements.

## Acknowledgements

MSB thanks Ministry of Education, Govt. of India for Prime Minister Research Fellowship (PMRF). MS thanks the Council of Scientific and Industrial Research (CSIR) for the fellowship. MAP thanks the University Grants Commission (UGC) of India for the research fellowship. SS thanks IISER Bhopal for the fellowship. The authors also thank the Sanger sequencing, NGS, and MASS facility at IISER Bhopal. The authors also thank the intramural research funds provided by IISER Bhopal.

## Authors contribution

VKS conceived and coordinated the project. MSB and MS collected the plant samples. MS performed DNA extraction, prepared the samples for sequencing, and performed Nanopore sequencing and species identification. MS and SS performed the RNA extraction from different tissues. MSB designed the computational framework. MSB performed the computational analyses presented in this study, analysed the data and constructed all main figures and tables. MAP performed the karyotype evolution and methylation analysis. MSB and VKS interpreted the results. MSB, MS, MAP and VKS wrote the manuscript. All the authors have read and approved the final version of the manuscript.

